# Sampling intensity and temporal persistence of airborne eDNA in partially enclosed spaces

**DOI:** 10.1101/2025.07.14.664745

**Authors:** Nina R Garrett, Orianne R Tournayre, Joanne E Littlefair, Natalia V. Ivanova, Guiying Mei, Tiffany Jedrecka, Andrew Briscoe, Amanda Naaum, Nancy B Simmons, Elizabeth Clare

## Abstract

Airborne environmental DNA (eDNA) has shown promise as a terrestrial biomonitoring tool and its ecological applications are expanding. Despite its growing use, airborne eDNA does not yet have the extensive body of supporting research like its aquatic counterpart, with considerable uncertainty remaining concerning how airborne eDNA behaves, with regards to signal duration, and how much sampling effort is needed to capture DNA in a given airspace. By using airborne eDNA in a semi-controlled environment which acted as an artificial roost where bat species and their abundances were known, we estimated the sampling intensity (both the number of samples and number of sampling events) required to capture bat diversity of a given airspace, as well as signal persistence of airborne eDNA. Together these data provide a temporal scale for airborne eDNA measurements. The majority of species richness was detected using as little as 4 samplers in this enclosed space and the greater the number of sampling events, the fewer samplers were needed. Both air movement and the type of environment (i.e., enclosed space, open area etc.) are likely to impact detection and need to be considered during study design. eDNA also appeared to settle out of the air quickly, suggesting that detections likely reflect recent activity, which also has important implication for rare species which may only have a narrow window for detection. Our results add to the growing body of literature that indicate airborne eDNA can be a useful biosurvey method, especially for rapid surveys in communities with high turnover rates.

## Introduction

In recent years, airborne environmental DNA (eDNA) has gained traction as a terrestrial biomonitoring tool (Johnson & Barnes, 2024). While the monitoring of pathogen transmission (e.g. Leung, 2021) has generated a growing body of literature, the aerial movement of other biological material in air, such as hair or skin cells from terrestrial mammals, has been less studied. Airborne material such as pollen (Craine et al., 2017; Mohanty et al., 2017) and fungal spores (Abrego et al., 2018) have been used to survey for local taxa, but on a limited scale. Johnson et al. (2019) tested for the presence of plant material in airborne dust and demonstrated that taxa that are not wind-pollinated could be detected as well as those reliant on wind dispersed pollen, indicating that biological particulates included non-pollen sources. Aalismail et al., (2021) found bird and mammal DNA in the global dust belt, and a proof-of-concept experiment demonstrated that terrestrial vertebrate eDNA could be directly retrieved from air (Clare et al., 2021). Since these discoveries, airborne eDNA has been used in a variety of studies ranging from detecting exotic species in zoos (Clare et al., 2022; Lynggaard et al., 2022) to surveying a plant community over a year (Johnson, Fokar, et al., 2021). Other demonstrations surveying for insects (Gregorič et al., 2022; Pumkaeo et al., 2021; Roger et al., 2022) and vertebrates (Johnson et al., 2023; Lynggaard et al., 2022; Sanchez et al., 2025; Serrao et al., 2021) have also emerged. In a first field application, airborne eDNA was used to survey bat populations in a series of Neotropical roosts (Garrett, Watkins, Francis, et al., 2023) including several judged too unstable for more traditional approaches like netting. More recently, airborne eDNA has been successful in detecting both threatened and invasive species (Frere et al., 2024; Geller & Partridge, 2025; Sanders et al., 2023). Geller and Partridge (2025) detected the invasive hemlock woolly adelgid (*Adelege tsuga*) in areas with no history of infestation. Doing so allowed for a more focused visual survey effort which led to the discovery of an infested tree on the site.

There are several benefits to the use of eDNA as a biodiversity survey tool. Because it does not require physical access to organisms, eDNA can be used to detect taxa that are cryptic or are no longer present in the immediate area, and, as a consequence, it may take less manpower to conduct a thorough survey (Thomsen & Willerslev, 2015). The methodology behind eDNA is less taxon specific, when compared with conventional biodiversity surveys, highlighting the potential to learn about many groups simultaneously. As in aquatic systems, airborne eDNA provides an alternative to the capture and physical handling of organisms (Garrett, Watkins, Francis, et al., 2023), which reduces the chances that researchers will disturb or disrupt the natural behaviour patterns they are studying. Airborne eDNA collection complements other existing survey and inventory methods, and can easily be deployed alongside stationary sampling devices such as camera traps (Polling et al., 2024). Airborne eDNA can be collected through passive sampling methods (Gregorič et al., 2022), allowing samplers to be deployed for days or weeks at a time (Johnson et al., 2023), or samples can be collected using active suction approaches (Clare et al., 2022; Lynggaard et al., 2022; Lynggaard, Frøslev, et al., 2023). An active approach allows for collection in areas without air flow (Garrett, Watkins, Francis, et al., 2023), and may increase collected yield/unit time.

Little is known about the temporal persistence of eDNA in air, and thus the time frame over which a detection may be informative has not yet been quantified (Johnson & Barnes, 2024). A few previous studies that have mentioned this issue reported contrasting results about the effect of sampling time, sampling duration and weather conditions on detectable biological material in the air (Després et al., 2012). In general, eDNA persistence is a function of creation, transport and degradation (Harrison et al., 2019). In aquatic systems, eDNA degradation is most often caused by microbial activity (Collins et al., 2018; Joseph et al., 2022). Particles can bind to sediment in aquatic systems, making them very stable and persistent. It is unknown if there is an analogous process in the air, but if something similar does occur it would allow some particles to persist longer than expected (Ferrer-Paris et al., 2013; Sakata et al., 2020; Turner et al., 2015). Wind can also resuspend eDNA particles, prolonging the detectable signal (Roger et al., 2022). Additionally, the behaviour of animals is often correlated with the environmental factors that may impact detection probabilities, making them difficult to disentangle. For example, rain can both wash particles out of the air and thus limit detection (Johnson et al., 2019, Milne et al., 2013) and increase natural activity patterns for certain species and thereby increase detection probabilities (e.g., the Texas toad (*Anaxyrus speciosus*) which only breeds in temporary pools formed after heavy rain, Johnson et al., 2023).

It is thought that most aquatic eDNA degrades quickly in the environment (3-7 days; Kucherenko et al., 2018; Williams et al., 2018) and only a small amount persists long term (Willerslev et al., 1999, 2003). As a consequence, we can hypothesize that only very abundant signals (i.e., those associated with very common species whose DNA is constantly being shed) would persist, though this is currently speculation with respect to airborne eDNA. In addition to degradation, it is unclear how quickly eDNA settles out of the air, although we can predict that settling rate will depend on particle size and local environmental conditions (e.g. wind and air currents). This creates uncertainty surrounding both the temporal persistence and degradation rates of airborne eDNA. Given these issues, it is unclear how much sampling is needed to capture the diversity of life in a given airspace - considerations that are particularly important when designing sampling strategies and interpreting the outcomes.

Unknown duration of signal persistence limits the ecological analyses and applications that can be conducted with airborne eDNA, particularly as compared to the common applications of aquatic DNA. Both marine (Jensen et al., 2022) and freshwater (Bista et al., 2017) community dynamics have been characterised with eDNA, but it is unclear at what scale airborne eDNA might be informative if used similarly. Johnson et al. (2023) found turnover in animals detected in airborne dust at two-week intervals, suggesting that eDNA signals will not persist over extended periods, and implying that airborne material could be a useful tool to monitor temporally co-occurring ecological activities. Garrett, Watkins, Francis, et al., (2023) noted a difference in day to night detections of bats and non-bat mammals, apparently reflecting use of the local area by these taxa.

In our previous experimental work (Garrett, Watkins, Simmons, et al., 2023), we deployed airborne eDNA samplers in a controlled “field lab” (a 5 x 7 m room originally built as a classroom) where bats of known species and abundance were being briefly held during a two-week annual bat research field trip in the Orange Walk district of Belize. In that trial, we recovered over 91% of the potential species richness (Garrett, Watkins, Simmons, et al., 2023). This site is ideal for the development of methodological protocols for three reasons. First, the bat community is complex and, due to field sampling onsite by many bat researchers, more than 1000 individual bats representing >30 species typically enter and exit the field lab during the normal two-week field research period. Second, because individual bats are brought to the field lab specifically for expert identification, the exact richness and abundance of the community is known along with the timing of entry and exit for any unusual captures. Finally, the room itself is a semi-controlled environment. For example, air moves freely through doors and windows, just as it would in a cave, or an artificial structure frequently surveyed for bats (e.g. a house or church), and ceiling fans keep air moving at a low and constant level, making sampling conditions more similar to the field than a completely closed environment.

Because the diversity of bats in the field lab reflects captures by field researchers sampling in nearby habitats, the room has a different community assemblage each night based on success of netting and trapping efforts by the field teams. As such this setting effectively acts as a temporary artificial bat roost site, but with a completely known temporal community composition. The field lab allows us to assess a bat community in an enclosed space, and because it starts as a novel environment empty of bat DNA, we can start to quantify the temporal persistence of airborne eDNA signals and assess the impacts of sampling intensity on data recovery. Given this experimental setting, our study had two interrelated goals: 1) to determine the eDNA sampling intensity (both the number of samples and number of sampling events) required to capture defined vertebrate diversity of a given airspace, and 2) to estimate the signal persistence of airborne eDNA in an enclosed environment, and to provide a temporal scale for airborne eDNA measurements.

## Methods

### Sample Collection

All work was completed under approvals from the York University ACC: 2021-10, American Museum of Natural History IACUC approval AMNHIACUC-20221110, Belize National Institute of Culture and History permit number IA/H/1/23/01, and Belize Forest Department permit number FD/WL/1/22 (54). Bats were captured using harp traps and mist nets in a variety of locations in and around Indian Church, Belize including two archeological reserves (Lamanai and Ka’kabish), sites on the New River (Barber Creek, Dawson Creek), and roosts in surrounding farmland. The landscape in the area consists of fragments of tropical dry forest (ranging from 450 ha to tiny patches) surrounded by an agricultural matrix. Bats were caught in the field and temporarily held in a series of clean cloth bags with each bag holding only one bat, and subsequently were brought back to the field lab (Figure 1) for identification and processing. The set of bats held in the lab changed each night reflecting the different sites where netting and trapping took place. The holding bags were washed in an industrial laundry between each use. Bats were briefly removed from the bags in the lab by experts in local taxonomy to confirm the species ID. After ID was confirmed, the bats either became part of other research occurring on site, which involved an increase in handling bats outside of the bat bags (e.g. collection of hair samples, PIT tagging), or they were removed from the field lab and were released at the point of capture. The length of time that any individual bat was out of the bag ranged from approximately 1 minute to 10 minutes.

**Figure 1.**
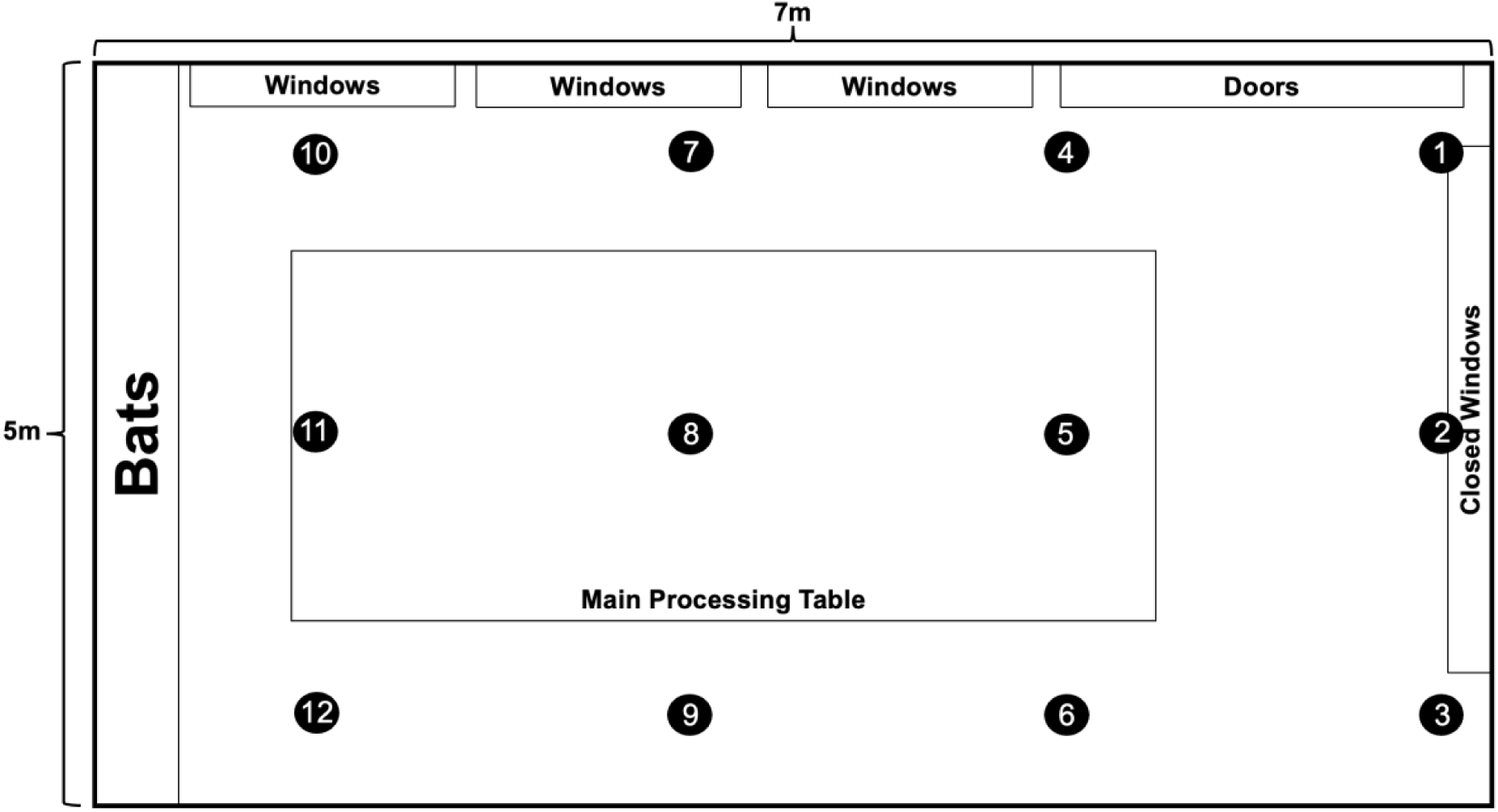
Sampler deployment grid (1-12). The room is a 5x7m rectangle with windows and a door along one long side and windows along one short side. The room is cement block construction with walls 3.6m high and a simple metal peaked roof. Two ceiling fans run most of the time, and windows and door are generally left open and are thus a source of external DNA.

### Air sampling

We deployed 12 purpose-built, 3D-printed 12V samplers (Garrett, Watkins, Simmons, et al., 2023, assembled following https://youtu.be/DHHR5Cg5dUE?si=DN1CUk_ADjX-qofT) in a 3x4 grid within the field lab (Figure 1). We cleaned each air sampler using a 10% solution of household bleach (6% sodium hypochlorite) and water before sampling and between every sample collection. Prior to field work, filters were cut from Filtrete 1900 furnace filters in a sterilized biological safety cabinet (BSC) and exposed to UV light for one hour before being placed in a sterile ziplock sandwich bag in sets of 12 for deployment. Samplers ran for 10 days and 10 nights for a total of 240 samples, each representing approximately 8 hours of sampling.

### Molecular Protocols

After air sampling, we removed each filter paper from the sampler and placed each filter paper in a clean ziplock bag and froze all the samples (-20°C) until they were moved to a molecular lab. Inside a biological safety cabinet (BSC), we cut a half circle out of the middle of each filter paper using scissors and forceps sterilized with a 10% household bleach solution and 70% ethanol between sampling each filter. We soaked each half circle in 2.5-3 ml PBS and incubated them at 50-56°C overnight in a rotating incubator. After incubation, we pipetted the PBS into a microcentrifuge tube and spun it down at 6000 x g for 3 minutes. We removed excess PBS leaving a small pellet at the bottom. In cases where a pellet was not visible, we left a small amount of PBS at the bottom to avoid removing the pellet. These steps were repeated until all the PBS had been spun down and material caught on the filter was concentrated into one microcentrifuge tube. We then treated the pellet as a tissue sample and DNA was extracted using the Qiagen Blood and Tissue kit (Qiagen Canada), following kit guidelines modified to incubate samples for 15 minutes in the ATL buffer and proteinase K and to use 100 µL of the elution buffer instead of the suggested 200 µL. Once the elution buffer was added, samples were incubated at room temperature for five minutes prior to being spun down.

To limit contamination, we extracted DNA in a BSC located in a controlled-entry room where activities are limited to only a few procedures. We cleaned the BSC and all equipment used to process samples (i.e., pipettes, racks etc.) with 1% Virkon, followed by 70% ethanol, and then exposed them to UV for 30 minutes before each use. After all lab processing was completed for the day, everything was again cleaned with 1% Virkon and 70% ethanol and exposed to UV for an hour.

We amplified all samples in triplicate via PCR using two mammalian targeted primers: Mam16s (∼90 bp) (Calvignac-Spencer et al., 2013; Taylor, 1996) and a bat specific COI primer (∼202 bp) (Walker et al., 2016). Our amplifications following the protocols in Garrett, Watkins, Francis, et al., (2023) were conducted with partners NatureMetrics in a dedicated clean room to limit contamination. Each 15 μL reaction mix included 7.5 µL of 2X QIAGEN Multiplex PCR Master Mix, 3 µL of ddH_2_O, 0.75 µL of each forward and reverse primer tagged with CS1 and CS2 adaptors (concentration 10 μM) and 3 µL of extracted DNA. For 16S our cycling conditions had an initial denaturation at 95°C for 10 min followed by 40 cycles of 12 s at 95°C, 20 s at 59°C and 25 s at 70°C and a final extension of 10 min at 72°C. For COI amplification our cycling conditions were 5 min for 95°C followed by 5 cycles of 1 min at 94°C, 1.5 min at 45°C and 1 min at 72°C followed by 35 cycles of 1 min at 94°C, 1.5 min at 60°C and 1 min at 72°C and then a final extension of 10 min at 95°C. We amplified DNA in 96 well plates and each plate included three negative controls (nuclease-free water), and three positive controls of *Cavia procellus* (Guinea pig) DNA as the templates, respectively. We visualized all PCR products on E-Gels (Invitrogen) and sent the products to the QMUL Genome Center in the UK. All samples, including positive and negative controls, were individually indexed, size selected using Ampure Beads (Beckman Coulter) and quantified and purified using DNA D100 TapeStation and QUBIT (Invitrogen). Sequencing was conducted on a MiSeq 2x300bp run with a MiSeq Reagent Kit v3. Raw sequences were demultiplexed on site..

### Bioinformatics

We processed all sequenced samples in the DADA2 (Callahan et al., 2016) pipeline in RStudio (RStudio Team, 2021). We used cutadapt (Martin, 2011) to remove primers, trim and filter sequences. Using the truncLen function, we removed low quality reads, and cut length was chosen for each marker based on approximate amplicon length and quality plots (16S = ∼90 bp, COI = ∼202 bp). The mergePairs and removeBimeraDenovo functions in DADA2 were used to merge reads and remove chimeras respectively. We manually checked each amplicon sequence variant (ASV) against GenBank records to determine an ID based on identity and query cover. IDs were only considered if they had 97% sequence identity and full query coverage with a reference of known provenance. These represent genus level matches. Species level matches were made with 100% sequence identity. Some names were manually curated where the existing reference retains old taxonomy (Appendix 1). ASVs not meeting these parameters were not included in further analysis. Before further analysis, all ASVs identified as human were removed along with other obvious sources of contamination such as aquatic species and bacteria. Each plate was then filtered based on the number of read counts in the negative controls associated with that plate (see discussion and Appendix 2 for a breakdown of negative controls). For the *Molossus* genus, we assigned ASV IDs only to genus level given the genetic similarity between species in this group at mitochondrial markers (Clare et al., 2011).

### Data Analysis Sampling Coverage

In this study, we refer to a sample as a single filter, a sampling event as a day or night where 12 samples were collected in a grid (Figure 1), and we also consider all samples collected over 10 days (n = 120 samples), 9 nights (n = 108 samples, May 2^nd^ night was removed because of contamination – see discussion), or the full set (n = 228 samples).

To determine the number of samples needed to detect the bat species richness in the room (i.e., sampling coverage) for both 16S and COI markers, we constructed an accumulation curve for each sampling event (i.e. a day or night that samplers were deployed; n_day_ = 10 and n_night_ = 9) using the R package iNEXT (Chao et al., 2014; Hsieh et al., 2024). To test for recovery efficiency, we calculated sampling thresholds (the number of samples needed to detect a certain proportion of species) of 85% and 95% taxonomic recovery following the methods used by Sellers et al. (2024). We also calculated the sampling coverage (the proportion of species richness detected in a given number of samples) after considering one, five, 10, and 20 samples (Table S1). We generated an overall accumulation curve for all 10 days sampled, for all 9 nights sampled, and one for the total sampling period. For these three curves, taxonomic coverage is described as measures of alpha diversity using Hill numbers which are equivalent to species richness (q = 0), the Shannon index (q = 1) and the Simpson index (q = 2) (Chao et al., 2014). For these curves, we generated an 84% confidence interval which equates to an α-level of 0.05 for overlapping distributions (Drinkwater et al., 2019; MacGregor-Fors & Payton, 2013) and extrapolated this to double the number of the observed values following Chao et al. (2014).

### Temporal Persistence

One of the largest unknowns in the use of airborne eDNA as a biodiversity survey tool is the temporal persistence of the signal. Little is known about whether a detection represents long term accumulation of eDNA which remains airborne, or if the signal is of short duration so that detections reflect only taxa most recently present nearby. We attempted to estimate signal persistence by concentrating on taxa that are rare within our community and hence the presence of one or more individuals at any time effectively constitutes a unique event. To determine the eDNA signal persistence in the enclosed space of our field lab, we tracked the detections of rare bat captures. For these purposes we considered every taxa for which we captured n = 1-3 individuals during the 10-day sampling period (Table 1). Read counts in each sequenced region and the days on which they were detected were recorded and compared to instances where the taxa were known to be physically present in the room and the sampling periods after. Doing so allowed us to establish an estimate of how long the DNA signal of an individual bat persisted once that individual was no longer present in the room. We anticipate that the eDNA signal from a super-abundant taxon will persist much longer because of the greater volume of biological material.

**Table 1.** Species and their abundance in the field lab during the sampling period and their record availability in the database for each marker. For each species present in the lab, the type of record in either the 16S or COI database is stated. A positive detection using eDNA is denoted with an “S” for species level detection and a “G” for genus level detection. “X” denotes no record or no eDNA detection. Discrepancies between the database and the eDNA detection are in bold.

| Genus | species | Abundance | 16S database record | eDNA | COI database record | eDNA |
| --- | --- | --- | --- | --- | --- | --- |
| *Artibeus* | *jamaicensis* | 41 | species | S | species | S |
| *Artibeus* | *lituratus* | 61 | species | S | species | S |
| *Artibeus* | *intermedius* | 26 | species | S | species | S |
| *Bauerus* | *dubiaquercus* | 1 | species | S | species | S |
| *Carollia* | *perspicillata* | 28 | X | X | species | S |
| *Carollia* | *sowelli* | 13 | species | S | species | S |
| *Dermanura* | *phaeotis* | 48 | species | S | species | S |
| *Desmodus* | *rotundus* | 92 | species | S | species | S |
| *Eptesicus* | *furinalis* | 35 | species | **G** | species | **G** |
| *Gardnerycteris* | *keenani* | 3 | X | X | species | S |
| *Glossophaga* | *commissarisi* | 7 | species | S | species | S |
| *Glossophaga* | *mutica* | 121 | X | X | species | X |
| *Lasiurus* | *ega* | 10 | species | S | species | S |
| *Lophostoma* | *nicaraguae* | 1 | species | S | species | S |
| *Lophostoma* | *evote* | 1 | species | S | species | S |
| *Micronycteris* | *microtis* | 1 | species | X | species | **G** |
| *Micronycteris* | *schmidtorum* | 3 | species | X | species | **G** |
| *Mimon* | *cozumelae* | 2 | X | X | species | S* |
| *Molossus* | *alvarezi* | 9 | genus | G | genus | G |
| *Molossus* | *nigricans* | 10 | genus | G | genus | G |
| *Myotis* | *elegans* | 19 | species | S | species | S |
| *Myotis* | *pilosatibialis* | 4 | X | X | X | X |
| *Natalus* | *mexicanus* | 4 | species | S | species | S |
| *Noctilio* | *leporinus* ** | 2 | species | **G** | species | S |
| *Platyrrhinus* | *helleri* *** | 3 | genus | G | species | S |
| *Pteronotus* | *fulvus* | 29 | species | S | species | S |
| *Pteronotus* | *mesoamericanus* | 81 | species | S | species | S |
| *Pteronotus* | *psilotis* | 2 | species | S | species | S |
| *Rhogeessa* | *aenea* | 15 | species | S | species | S |
| *Rhynchonycteris* | *naso* | 19 | species | S | species | S |
| *Saccopteryx* | *bilineata* | 21 | species | S | species | S |
| *Sturnira* | *parvidens* | 54 | species | S | species | S |
| *Trachops* | *coffini* | 1 | species | S | species | X |
| *Uroderma* | *convexum* | 24 | species | S | species | S |
| *Vampyressa* | *thyone* *** | 2 | genus | G | species | S |
| # of Detectable Species |  |  | 26 |  | 32 |  |
| # of Detectable Genus |  |  | 4 |  | 2 |  |
| Total Detectable |  |  | 30 |  | 34 |  |
| Total Detected using eDNA |  |  | 28 |  | 32 |  |

\*This detection was below the negative control filtering threshold and removed from further analysis
\*\* *Noctilio leporinus* was not detected despite a species level record in the database, however the sister species *N. albiventris* was detected. Given that *N. albiventris* is not present in the study area, this detection was treated as *N. leporinus*.
\*\*\* There are no species-level records for *Vampyressa* and *Platyrrhinus* using the 16S marker in the database for the species present at this field site but for both only one taxa is known from the area so these detections are reported to species level rather than genus

## Results

After merging and chimera removal, we recovered 7.7 million reads representing 865 ASVs for the 16S marker. After removing ASVs that did not meet filtering requirements (i.e., matches lower than 97%, less than 100% overlap, human or other obvious contamination matches), 189 ASVs remained in the 16S data. For the COI marker, we recovered 6.6 million reads representing 1308 ASVs. After filtering, 724 ASVs remained in the CO1 data. With the exception of plate seven in the 16S data, and one species match in plate 2 of the COI data, negative contamination was minimal (<36 reads) (Appendix 2). There was a total of 35 species present in the field lab during the two-week sampling period, but not all species had records in the Genbank database (Table 1). Of the species represented in reference collections, we detected 93% of the species using the 16S region and 94% using the COI region.

During our study 30 taxa contained references for the 16S marker and using these we had the potential to identify 26 to species level and 4 to genus level (Table 1). Using airborne eDNA, we differentiated 22 taxa to the species level, and six to genus. Of the latter six taxa, two only had genus level records in the database. Two were in the database (*Molossus alvarezi* and *M. nigricans*) but cannot be distinguished reliably using the 16S marker and the short DNA fragments used in our study, so we kept these identified to only the genus level. The other two sequences are in the database, but we could not reliably match them at the species level. In the case of *Noctilio*, there is only one species known to be present in our research area, we but we retained it identified to only the genus level here to be conservative. Only two “detectable species” using the 16S marker, *Micronycteris microtis, and M. schmidtorum,* were not found among our eDNA samples despite having been captured and processed in the field lab; however, both species were among the rarest taxa (≤ 3 individuals captured) in our study.

In comparison to the 16S marker, 34 taxa were potentially identifiable using the COI marker, 32 of which could be identified to species level and two to genus level. Of these, we identified 27 taxa to species level and five to the genus level. Two of the detections (*Molossus alvarezi* and *M. nigricans*) at the genus level are missing species records in GenBank (Table 1). The other three are in the database but we could not reliably differentiate them below the genus level. Within the COI data, we did not detect *Trachops coffini* (rare) or *G. commissarisi* (also relatively rare, only 8 individuals captured).

### Sampling Coverage

To quantity the effectiveness of our sampling strategy on any given sampling event (one day or one night), we compared the number of samples needed to reach a threshold of 85% or 95% coverage of known bat species present in the room. For the 16S data, the mean sampling threshold was 3.7 samples for the 85% threshold and 8.7 samples for the 95% threshold (Figure 2A). For the COI region, the same detection coverage was reached with 2.8 (85% threshold) or 7.4 (95% threshold) samples in any given event (Figure 2B). Any one sample taken at random captured at least 40% of richness in the room using either marker (Table S1). For each sampling event, we detected most of the taxa present, with most sampling events missing fewer than two species known to be present at the time (Figure 3A).

**Figure 2.**
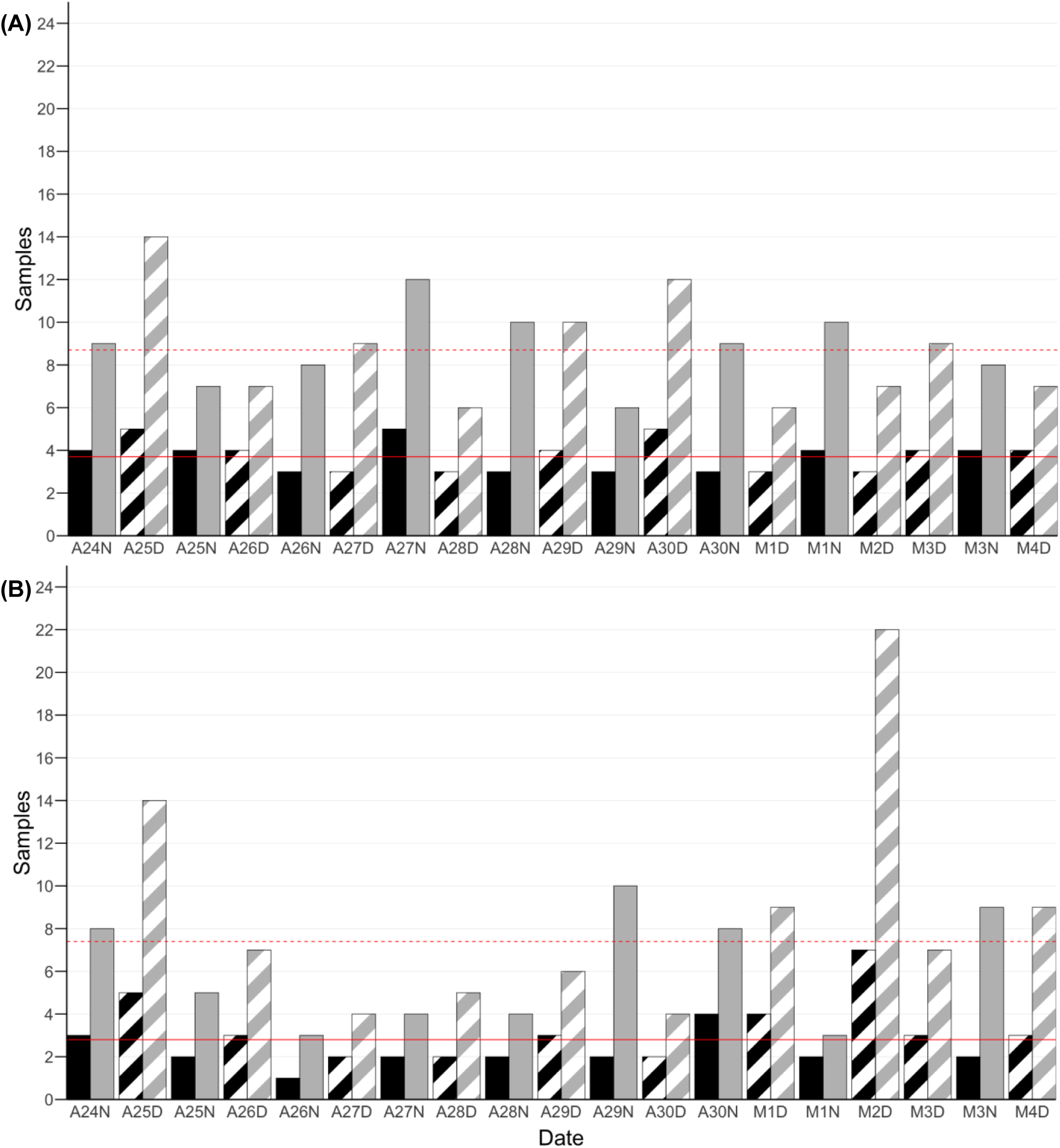
The number of samples needed to detect 85% (black) and 95% (grey) of the species in the room for each sampling event (night: solid, day: striped, e.g., A24N = April 24^th^ night, M1D = May 1^st^ day) using the 16S (A) and COI (B) regions. The average number of samples for each sampling threshold are indicated by a solid red (85%) and dashed red (95%) line.

**Figure 3.**
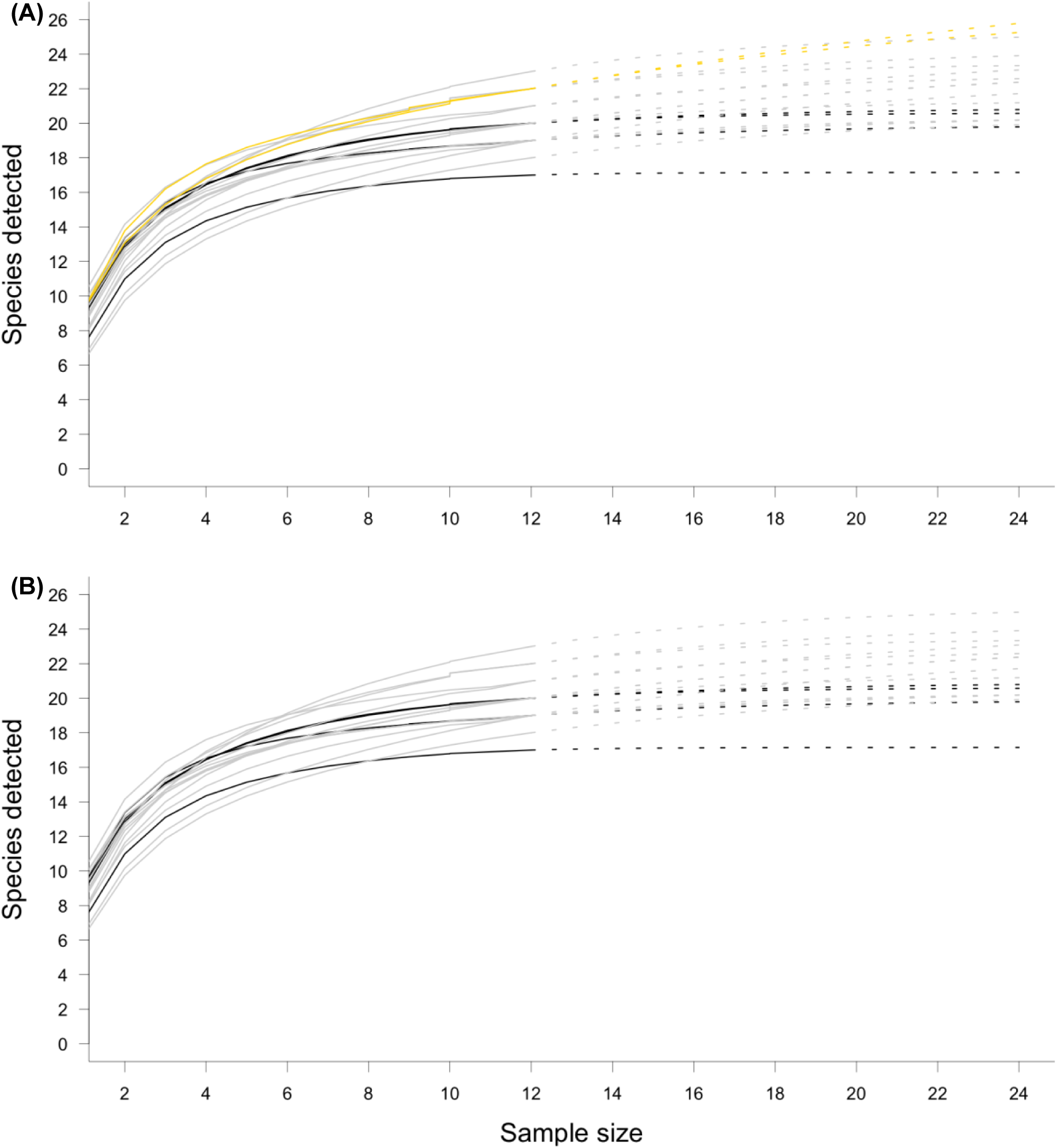
Accumulation curves for bat diversity detected during each sampling event using the 16S (A) and COI (B) markers. For each region, the number of undetected species for each sampling event are indicated as follows: less than one undetected species is indicated with a black accumulation curve, less than 2 undetected species in grey and less than 4 undetected species in yellow. Estimates have been extrapolated to 2x the sampling.

For the entire sampling period, taxon accumulation curves reached an asymptote quickly for all three estimated parameters (q = 0 - 2) in 16S and for q = 1 (the Shannon index) and 2 (the Simpson index) in the COI marker in all instances (10 days, 9 nights or the full set of 19 events; Figure S1). Because the community is dominated by common taxa, 85% of the total species richness could be detected with four random samples despite variation in the community from day to day.

### Temporal Persistence

Of the 35 bat species which were physically brought into the field lab, 10 were caught on three or fewer occasions, and hence represent “rare” species whose DNA would not be expected to be continually present (Table 1). We tracked 17 captures of rare species to assess the persistence of the eDNA signal.

In eight cases, eDNA detected a rare species only on the night that it was present in the field lab (e.g. *Lophostoma evote*, April 30 night, COI) (Table 2). In seven cases, rare taxa present in the room were not detected with airborne eDNA the day of their capture nor the following day (e.g. *Micronyteris sp*., April 25 night, COI) (Table 2). In two cases, taxa were not detected with eDNA on the night of capture but were detected in a subsequent event (e.g. *Mimon cozumelae* COI, *Bauerus dubiaquercus* 16S). In 13 cases, the signal persisted across multiple sampling events, with consistent (e.g. 3 sampling events in a row for *Gardnerycteris keenani* COI) or fluctuating (e.g. *Lophostoma nicaraguae* COI) eDNA detections over time (Table 2). Only *Noctilio sp.* (both markers; Table 2) (see discussion) was detected prior to the night it was physically present in the room. The only species missed entirely was *Trachops coffini* using the COI marker.

**Table 2.** Read counts by sampling event for each of the rare species caught ≤ 3 times. Where relevant, the total number of bats in the room are indicated in brackets. Green shading represents when taxon were present in the room. Black indicates cases where the taxon was detected with airborne eDNA only when present in the room, grey indicates eDNA detection on the night of capture and subsequent days, and blue indicates an absence of eDNA detection on the night of capture but positive eDNA detection in the subsequent events. Cases where taxon was present but no detected have no box. Detections are split by marker. There are four cases in the 16S region where a species level detection could not be made and thus, were recorded at the genus level, indicated by purple text.

|  | COI |  |  |  |  |  |  |  |  |  | 16S |  |  |  |  |  |  |  |
| --- | --- | --- | --- | --- | --- | --- | --- | --- | --- | --- | --- | --- | --- | --- | --- | --- | --- | --- |
|  | *Bauerus dubiaquercus* | *Gardnerycteris keenani* | *Lophostoma evotis* | *L. nicaraguae* | *Micronycteris* sp. ** | *Mimon cozumelae* | *Noctilio* sp. | *Platyrrhinus helleri* | *Trachops coffini* | *Vampyressa thuyone* | *Bauerus dubiaquercus* | *Lophostoma evotis* | *L. nicaraguae* | *Micronycteris* sp. ** | *Noctilio* sp. | *Platyrrhinus helleri* | *Trachops coffini* | *Vampyressa thuyone* |
| April 24 Night (n = 26) | 4 | 0 | 0 | 0 | 0 | 0 | 0 | 0 | 0 | 6732 | 0 | 0 | 0 | 0 | 0 | 57 | 0 | 654 |
| April 25 Day | 6 | 0 | 0 | 0 | 0 | 0 | 0 | 0 | 0 | 0 | 0 | 0 | 0 | 0 | 0 | 162 | 0 | 0 |
| April 25 Night (n = 150) | 210 | 0 | 6 | 0 | 0 | 0* | 0 | 378 | 0 | 840 | 9 | 11 | 0 | 128 | 0 | 939 | 151 | 29 |
| April 26 Day | 3 | 0 | 0 | 0 | 0 | 0 | 0 | 0 | 0 | 245 | 0 | 119 | 0 | 0 | 121 | 0 | 170 | 0 |
| April 26 Night (n = 50) | 0 | 5876 | 0 | 73 | 0 | 4 | 320 | 0 | 0 | 0 | 0 | 7 | 226 | 0 | 0 | 0 | 0 | 33 |
| April 27 Day | 0 | 43 | 0 | 0 | 0 | 0 | 9 | 0 | 0 | 0 | 0 | 46 | 0 | 0 | 0 | 0 | 0 | 0 |
| April 27 Night (n = 82) | 0 | 2257 | 0 | 222 | 0 | 0 | 0 | 121 | 0 | 0 | 0 | 84 | 131 | 0 | 0 | 248 | 42 | 0 |
| April 28 Day | 0 | 134 | 0 | 212 | 0 | 0 | 0 | 0 | 0 | 0 | 0 | 0 | 138 | 27 | 0 | 16 | 0 | 0 |
| April 28 Night (n = 111) | 0 | 2147 | 0 | 0 | 0 | 0 | 0 | 0 | 0 | 0 | 0 | 0 | 0 | 29 | 0 | 7 | 0 | 0 |
| April 29 Day | 14 | 3 | 0 | 0 | 0 | 0 | 0 | 0 | 0 | 0 | 0 | 0 | 0 | 0 | 0 | 0 | 15 | 0 |
| April 29 Night (n = 31) | 457 | 0 | 0 | 0 | 0 | 0 | 0 | 0 | 0 | 0 | 0 | 0 | 0 | 0 | 0 | 0 | 0 | 0 |
| April 30 Day | 186951 | 0 | 0 | 0 | 0 | 0 | 7 | 0 | 0 | 0 | 7742 | 0 | 0 | 0 | 0 | 0 | 110 | 0 |
| April 30 Night (n = 41) | 41941 | 0 | 1201 | 0 | 0 | 0 | 11 | 0 | 0 | 0 | 243 | 1414 | 0 | 0 | 0 | 835 | 0 | 0 |
| May 1 Day | 2544 | 0 | 0 | 0 | 0 | 0 | 2 | 0 | 0 | 0 | 21 | 0 | 0 | 0 | 0 | 0 | 0 | 0 |
| May 1 Night (n = 85) | 5510 | 0 | 0 | 0 | 22 | 0 | 0 | 0 | 0 | 197 | 12 | 78 | 0 | 28 | 0 | 0 | 0 | 0 |
| May 2 Day | 0 | 0 | 0 | 0 | 0 | 0 | 0 | 0 | 0 | 0 | 7 | 0 | 0 | 0 | 0 | 0 | 0 | 0 |
| May 3 Day | 7 | 0 | 0 | 0 | 0 | 0 | 0 | 0 | 0 | 0 | 0 | 0 | 0 | 0 | 0 | 0 | 0 | 0 |
| May 3 Night (n = 61) | 12 | 372 | 0 | 0 | 0 | 0 | 0* | 0 | 0 | 0 | 0 | 0 | 628 | 0 | 2903* | 0 | 0 | 0 |
| May 4 Day | 0 | 0 | 0 | 0 | 0 | 0 | 0 | 0 | 0 | 0 | 0 | 0 | 0 | 0 | 0 | 0 | 0 | 0 |

\* More than 1 individual present
\*\* These detections are of a *Micronycteris* species not found in the area. The match only had a 94% similarity and thus was kept to genus level. It was retained here as a low-quality detection. Note that May 2 night was removed, see discussion.

## Discussion

Our objectives were to determine the sampling intensity required to capture the diversity of mixed bat community in a roost (e.g. cave, building), and to estimate the period of time for which a novel airborne eDNA signal would remain detectable using our inexpensive purpose build samplers. Our goal is to provide experimental evidence for how to maximize sampling and provide a temporal scale for detections of rare species of interest. In general, our data suggests that surveying a small air space such as a building or other small bat roost (e.g., cave or hollow tree) does not require extensive sampling to recover most of the diversity of taxa present. In our experiment, 3-4 samplers running for a few days and nights recovered >85% of the taxa present in the space. Doubling this to 7-8 samplers could recover almost all the diversity in the room in a single night, suggesting a simple trade-off between sampling intensity and duration, with a small variance based on which genetic markers are used and the completeness of a reference database.

Our data suggest that airborne sampling can be used as a very effective mass sampling system but there are several factors that need to be considered when planning the sampling effort to be employed. In particular, the type of community being surveyed (i.e., a community with day-to-day variability vs. consistent composition) and the focus of the study (i.e., common species or rare species) may dictate the sampling intensity. This is particularly true when we consider results of our analysis of the signal persistence of rare taxa, which suggests that eDNA signals from such species may only last a matter of hours or a day or two, and that detections reflect only very recent activity in an area. As such, a greater sampling effort over a longer period of time would be required to detect relatively rare species, as is the case with most sampling methods. When communities have high day-to-day turnover, sampling over more days is likely to be more useful for assessing overall species composition of the community than intensive sampling on a single night since eDNA may settle out of the air and become quickly undetectable. Accordingly, the signal from a rare taxon or one only occasionally present in a roost would not be expected to persist long-term, and detection of that species in the community may depend on sampling either at the time that the animal is present or soon after.

### Documenting the dynamics an ecological community

Two different objectives are common in community sampling: (1) rapid assessment of community membership (a simple survey of what species are present), and (2) longer term assessment of community dynamics (e.g., changes over time). The optimal airborne eDNA sampling approach will change depending on which objective is the focus of the study. In cases where rapid screening is the goal, airborne eDNA assessments of multiple locations simultaneously would likely require less survey effort. For example, surveying many buildings for the potential presence of bats (common in some environmental consulting applications) may require only one night of sampling as a screening approach. Using a grid of 12V samplers, we were able to successfully detect almost all taxa in the room during any given sampling event. Alternatively, longer term analyses may be used to assess community dynamics and search for rare taxa. For example, studies of roost use and community turnover may require longer term monitoring than other types of surveys. Klepke et al., (2022) found a higher species and ASV richness the longer samples were exposed to the air. Likewise, when we consider the full sampling period (ten days and nine nights), just four of the 228 samples taken in our study were enough to detect 80% of species also suggesting that a longer sampling period results in a higher richness recovery but with rapidly diminishing returns. On any given day or night sampling event, four samplers running in parallel would achieve a similar success rate with some variance depending on the marker considered.

Our results suggest that all common species (Shannon and Simpson estimates; Figure S1) and most rare species are detectable with much less sampling effort than we employed during our study. This is not surprising since the ability to detect the majority of taxonomic diversity in a few samples has also been documented in aquatic eDNA research (Beentjes et al., 2019). Our data also provide evidence that sampling in a grid-like pattern is an appropriate strategy for enclosed spaces such as small caves and buildings when the goal is to gain a general knowledge of use of the space overall. It should be noted that the situation in our field lab would be considered a situation with very high turnover involving low abundance and high diversity. This kind of situation presents a particular challenge for detecting rare taxa, but it avoids the potential problem of one superabundant species generating such a strong eDNA signal that it masks or reduces detection probability for a rare individual of another taxon.

Some contextual factors are key in designing a sampling scheme for a particular project and one of these is the timing of deploying samplers. For example, when comparing any given nighttime sampling event in our study with detections during the following daytime period, we found that sampling estimates tended to be lower during the day sampling events, and/or require greater sampling intensity (Figure 2) to achieve the same coverage. This reflects the fact that the room is mostly empty of bats during daytime sampling periods – a case of a true negative rather than undocumented diversity. Observed drops in recovery rate and estimates of the need for increased sampling intensity are indicative of short signal persistence for eDNA in the room. In designing eDNA studies and analyzing results, it is important to differentiate increased sampling intensity (more samplers run simultaneously) from increases in the number of sampling events (sampling on different days or at different times). In this study, where there was variability in the species and numbers of bats present during each sampling event, collecting fewer samples during any given event but spreading sampling events over a longer time period was shown to be key to capturing the total diversity of species present, particularly given the short duration of the eDNA signal.

The sampling coverage required to detect the full diversity of a community is likely to differ in a fully open space compared to an enclosed space given differences in air movement. When sampling aquatic eDNA in isolated dune lakes, Beentjes et al. (2019) found that there were large differences in samples both spatially and temporally, leading us to expect that in natural airspaces there will be more spatial heterogeneity than in enclosed rooms or other small spaces like caves. Various environmental factors including air flow patterns can also affect eDNA signal distribution (Altermatt et al., 2023). For example, the ceiling fans in our field lab likely caused a degree of air mixing, leading to more homogeneity of signal distribution and decreasing temporal variability, things that would not necessarily be the case in a natural bat roost unless there was air flow or the animals were frequently moving (which might be the case in many natural bat roosts). Considerations of air movement patterns might suggest a very different approach to eDNA sampling design based on site use, e.g. at a swarming site (Fenton, 1969) where bats are in a constant state of high activity. In the case of a bat swarming site, sampling for a shorter period of time with fewer samplers in a uniform distribution would probably capture sufficient diversity since movement in the roost is expected to result in continuous DNA shedding and a relatively homogenized signal. In a hibernaculum where bat activity is minimal and air homogenization is limited, sampling for longer periods with more samplers close to where the bats are roosting will likely have better success at detecting diversity.

In our study, we failed to detect *Micronycteris* and *Glossophaga commissarisi* despite these taxa being present in the field lab. While in our case we can distinguish a true negative from a false negative given that we know definitively what species were in the room, biosurvey methods in general face the challenge of distinguishing true from false negatives when deployed in areas where faunal composition is unknown or only partially known. Factors that make a taxon more “detectable” by airborne eDNA are as yet unknown, though there was a relationship in our data with rarity. The problem of “missing” taxa or false negatives is common with all survey methods (Tulloch et al., 2025). In previous airborne eDNA experiments centered on zoo systems with known assemblage membership, Lynggaard et al. (2022) did not detect wallaby (*Macropus rufogriseus)* and Clare et al. (2022) noted the absence of the Maned wolf (*Chrysocyon brachyurus*) from inventories despite these animals being present in the area at the time of sampling. False negatives have also been observed in aquatic studies (e.g., no strong positive detections for *Orcinus orca* despite sampling within 20 m of them (Pinfield et al., 2019).

In contrast to false negatives, we detected eDNA from *Noctilio* in the lab before it was actually brought into the space. It is likely that these detections resulted of secondary transfer from equipment used to in a study of *Rhynchonycteris naso*, an emballonurid frequently captured but not brought back to the lab. *Noctilio* and *Rhynchonycteris* both forage over water and are caught using nets strung from a boat. Often *Noctilio* hit the nets but escape before becoming entangled. The nets and other equipment used on the boat are stored in the field lab and moved every evening and morning, activity that may have resulted in the introduction of the DNA signal from *Noctilio* before any of these bats were brought into the lab. This type of contamination is a problem that must be accounted for in studies where field equipment is used repeatedly in different settings.

### The temporal window for detections

Based on our results, individual airborne eDNA signals are temporally localized with species generally only being detected within a 24-hour window while they were present and just after. In a few cases signals from rare taxa apparently lasted as long as 72 hours and became detectable again later on despite the animal not being present – or detected via eDNA – in the intervening time. The most likely explanation for these observations is the resuspension of eDNA due to disturbance in the room. The field lab is a very busy work environment and as people move equipment (i.e., gloves, calipers, bat bags, boxes, etc.) DNA almost certainly becomes re-airborne. Environments subject to less disturbance, such as a natural bat roost in a cave or hollow tree, would likely produce fewer secondary spikes in DNA.

The short timeframe that we observed for persistence of airborne bat eDNA in our field lab has yet to be reflected consistently in other eDNA research. In general, such short-term signals could be used to track rapid community composition changes. In airborne eDNA, Lynggaard et al., (2024) detected a shift in the vertebrate community over time but reported no clear effect of day vs night sampling, suggesting eDNA persists longer than 12h. While the amount of DNA detected tends to decrease rapidly (Strickler et al., 2015), eDNA in freshwater can be detectable anywhere from two days (Seymour et al., 2018) to 58 days after initial detection (Strickler et al., 2015). In marine environments, DNA can be detected from 14 to 26 days after the initial detection (Kutti et al., 2020; Weltz et al., 2017). In both cases, temperature and pH are thought to influence temporal persistence (Breton et al., 2022; Collins et al., 2018; Kutti et al., 2020; Strickler et al., 2015; Weltz et al., 2017). Our shorter time frame for airborne eDNA persistence may reflect material settling out onto surfaces (and hence unavailable to our samplers) rather than degradation, and because we focused only on rare taxa our estimates represent the persistence of an individual signal. The short-term nature of an individual airborne eDNA signal from a rare species could provide evidence that a site has been used or occupied very recently by the target organism, which also means that its detection window would be small. The persistence of an individual signal is particularly useful as an estimate for use in studies detecting at-risk species but also suggests that a greater need for dense, repeated sampling to maximize the likelihood of detection.

### Non-target detections and contaminant tracing

Because airborne eDNA analysis is a developing method, it is important to consider how detections are made and potential sources of contamination. While we saw little evidence of contamination, a full contaminant tracing analysis was completed (Appendix 2). A few key findings are considered here. First, we observed spikes in read counts that we consider to be false positive detections (identifications with high percentage identity and query coverage when the animal was not present) including *Lophostoma nicaraguae* (present April 26), *Gardnerycteris keenai* (present April 28) and *Noctilio* (never present in the field lab prior to May 3^rd^) on May 2^nd^, and sequence read counts from that night are an order of magnitude higher than other sampling events. We excluded cross contamination or batch effects during PCR because these samples were in the same batch as those the day of May 3^rd^, where we did not see the same effect. We also excluded contamination during DNA extraction and PCR as the relevant negative controls had only 2 reads, making contamination unlikely. The most likely explanation for the pattern observed is contamination during field sampling on May 2^nd^. We suspect two potential sources. May 1^st^ and 2^nd^ are the days when part of our field team packed up and left and new team members moved into the field lab, a process that involves a lot of movement of equipment, boxes, and duffle bags in the lab. These days would be the most likely days for high dust disturbance, and a week’s worth of settled biological material may have been resuspended quite suddenly. The second observation of interest is that Sampler 4 was located near the main garbage bag where consumables are deposited each night. A secondary garbage bag is often located near sampler 6, and sampler 5 is between them in a line (Figure 1). Though we did not record this, it is likely that garbage bag was changed when new team members arrived May 1^st^ and 2^nd^. All the detections in these three samplers showed increased read counts for this sampling event. We strongly suspect that the garbage acted as a “reservoir” of DNA that became a source when disturbed. While we cannot confirm this scenario, we treated May 2^nd^ detections with caution and removed them from our analysis (thus we consider only 9 nights in subsequent analyses). It is also an important reminder of the importance of documenting field activities and field environment, which in this case allowed us to perform contaminant tracking and generate a reasonable hypothesis for the unusual results for this sampling event.

A continual challenge specific to airborne eDNA analysis is to develop effective methods for establishing field controls. In aquatic systems, it is customary to run sterile water through sampling equipment at regular intervals to quantify contamination and ensure that field sterilization procedures are operating. No equivalent has yet been developed for sampling air (Tulloch et al., 2025). Our best option appears to be documenting what else is happening in the environment (for example, the position of garbage receptacles in our room). This allowed us to exclude one sampling instance which appeared to have been contaminated in the field. While not as quantifiable as sterile water in aquatic eDNA, it was justifiable in our case.

We observed minimal evidence of molecular lab-based contamination after the removal of ASVs considered ubiquitous (e.g. human) (Appendix 2). The exception was plate seven, where >1000 16S reads from two bat species were found in one negative control of one replicate. Because this was seen only in one of many controls, we did not use full negative filtering but excluded detections of species that did not have read counts higher than the negatives within that plate. Filtering criteria that are too strict can result in the loss of large portions of usable data and inflate false negatives (Garrett, Watkins, Simmons, et al., 2023). Altering the filtering criteria specifically for plate seven is a reasonable and informed way of using complex replicate, batch and control designs as a guide to retain as much data as possible while being stringent in determining true positives. For the COI region, the highest bat contaminant in the PCR negative controls was 233 reads of a *Molossus sp.* (Appendix 2). With low read counts in the negatives for all the plates amplified using the COI region, normal negative control filtering was used.

In addition to false positive and false negative detections, there are also false negative and false positive identifications, where an ASV has a real biological origin during sampling, but its identity cannot easily be determined. Particularly in Central America, bat taxonomy has been in considerable flux (Tulloch et al., 2025). Among the 50 species known from this field site, there have been at least 21 recent nomenclature changes (Appendix 1). To further complicate this, most old names remain valid in other areas of the species’ range so the same name attached to two different records in Genbank may be correct in one case and incorrect in another, and which is which can be clarified only when geographic ranges and taxonomic concepts are considered. Changing taxonomy alone creates considerable confusion when using public databases as a source for eDNA identifications, and such issues can inflate richness estimates if care is not taken. When the validity of a name is geographically linked the complexity increases, and autoclassification systems can generate both false positive and false negative errors. In addition to name changes, databases are limited to the records they contain. If DNA sequences for a species have not been reported for 16S or COI markers, those records will not be in the database. This can generate false negatives since any ASV associated with that species will generate no match despite the species being present at the site and its DNA in your sample.

It is only because our research team includes several experts in the taxonomy of the bat populations at our study site that we can correct the nomenclature for the purposes of our own reporting. In example, an auto-classifier might have recorded the presence of both *Carollia brevicauda* and *Carollia sowelli*, the first being a name now only valid in South America but both commonly attached to records in from Central America. Without correction, our species count would have increased in error. To the best of our knowledge, the taxonomic names used here are correct at the time of publication although many of the reference sequences we used are archived under out-of-date taxonomic concepts as in the example above, and we anticipate more nomenclature revisions in the near future. We went to considerable effort to trace the provenance of each reference sequence in order to validate and correct the references to the taxonomy used here. Despite this effort, there are still a few cases of identifications for which we cannot yet confirm the source of the reference sequence(s), and these were excluded from analysis. Caution should be exercised when using public databases as references in eDNA studies. This applies to all metabarcoding and eDNA work where it is impossible to trace taxonomic changes, particularly when an analysis contains thousands of taxa across dozens of orders and phyla. Taxonomic identifications in eDNA studies should be treated as hypotheses, and they are only as good as the databases they are built upon.

## Conclusions and future considerations

The results presented here provide a first estimate of sampling effort and airborne eDNA turnover rate in a semi-enclosed environment under partially controlled conditions. Based on these results, we conclude that if the aim of airborne eDNA sampling is to determine species evenness (the Simpson index) or both evenness and abundance (the Shannon index), deploying a few samplers is likely sufficient even in a complex community. If, however, the aim is to capture total species richness, more samplers are likely necessary. If the goal is to detect a rare species, increasing the number of sampling events is also likely required. Our data also indicates that DNA from an individual settles out of the air relatively quickly, in some cases in a matter of hours, reflecting recent activity. However, it is currently not clear how these estimates might have changed if the space has been isolated without any air flow or disturbance, and we anticipate that signals from super abundant taxa will persist much longer.

While we attempted to simulate real environments where the use of airborne eDNA sampling is being proposed (e.g. the survey of buildings/churches), more work is needed to test temporal persistence in an open area or natural roosts. We also need a much better understanding of the relationship between particle size and signal persistence. In this study, we use simple collectors which sample a larger range of particle sizes, as these types of samplers are the most likely to be used in field studies where costs need to be kept low, and ease of transport and repair are paramount. These home-made samplers are simple, easy to make and inexpensive. Commercially available air samplers are currently excessively expensive for this application but may have better ability to restrict particle sampling size which could impact signal persistence. A better understanding of the dynamic between persistence and particle size could help researchers make better informed decisions regarding equipment and study design. The short duration of signals measured here suggest that airborne eDNA sampling can be a very useful tool when the goal is rapid surveys of dynamic communities with suspected high turnover. Our results suggest that airborne eDNA can be a useful survey tool which reflects recent activity. While there are still unknowns that could limit its application, it can help pinpoint areas that require more intense surveying with a boarder range of methods and can help direct researchers to specific areas of interest particularly when resources are limited. Its ability to survey several locations at once with minimal manpower and the low cost associated with the sampling method used in this study highlights the practical value of this method for biodiversity monitoring.

