## Supplementary material for "Sampling intensity and temporal persistence of airborne eDNA in partially enclosed spaces": Suplement File - Appendices

**Supplemental File: Appendices 1-4**

**Appendix 1:** Recent taxonomic revisions have altered the names of many bat taxa in our study area. These names still populate reference libraries, and many are valid in other parts of the range generating a significant taxonomic challenge. For example, *Artibeus* is still a valid genus, but all smaller bodied taxa are in the new genus *Dermanura*. Thus, records in Genbank exist under both generic names. Similarly, *Carollia brevicauda* used to be the name applied to all individuals. The name is now valid in South America while *C. sowellii* is for Central American individuals. However valid records from Central America are found in collections with both names attached.

In this list the older names are given first, following by the name changes over the period of work at this site with the name accepted for the taxa at the time of sample collection.

(small) *Artibeus = Dermanura*
*Tonatia = Lophostoma*
*Lophostoma brasiliense = Lophostoma nicaraguae*
*Uroderma bilobatum = Uroderma convexum*
*Carollia brevicauda = Carollia sowelli*
*Sturnira lilium = Sturnira parvidens*
*Glossophaga soricina = Glossophaga mutica*
*Mimon bennettii = Mimon cozumelae*
*Mimon crenulatum = Gardnernycteris crenulatum = Gardnernycteris keenani*
*Pteronotus parnellii = Pteronotus mesoamericanus*
*Pteronotus davyi = Pteronotus fulvus*
*Pteronotus personatus = Pteronotus psilotus*
*Myotis keyaysi = Myotis pilosatibialis*
*Lasiurus blossevilli = Lasiurus frantzi*
*Molossus ater = Molossus rufus = Molossus nigricans*
*Molossus sinaloe = Molossus alvarezi*
*Natalus strameneus = Natalus mexicanus*
*Rhogeesa aeneus = Rhogeesa aenea* (a spelling change)

**Appendix 2:** Contaminant tracing

**Negative lab controls in 16s data.**

Of the 10 plates, four plates (1, 4, 6 & 8) amplified with the 16S markers yielded no sequence data in the negative controls (Table S1). In plate 2, only one of the negatives yielded sequence data with 2 reads from a single ASV. This ASV was a 100% match to both *Artibeus intermedus* and *Artibeus literatus*. Both of these species were present in the room and met our filtering criteria (100% match). Plate 3 also yielded no sequence data in two of the three negative controls. A single control had sequence data from two ASVs, one with a 100% match to *Desmodus rotundus* (8 reads) and a 93% match to *Alouatta palliata* (2 reads) (Table S1). Given its lower match percentage and is not a bat species, *Alouatta palliata* was discarded. As in plate 2, *D. rotundus* met our filtering criteria and was present in the room. Plate 5 had two negatives that yielded sequence data. Two of the ASVs found in theses controls were the same as those present in plate 2 and 3 (matched to *A. intermedius/literatus*: 3 and 2 reads and *D. rotundus*: 5 and 3 reads). The third ASV was a 100% match to *Pteronotus mesoamericanus* (3 reads) which was present in the room. The final ASV was a 100% match with *Cavia porcellus* (18 reads in one control) which was the species used as the positive control in all plates (Table S1). Thus, this detection was removed. While plate 7 had the highest read counts in the negative controls, it was limited to just one of the controls. Sequence data in this control matched to 6 ASVs. Two were to a 100% and 98.89% match to *Carollia perspicillata*. Only the ASV with a 100% match met filtering criteria and was retained. It yielded 1430 reads. The only other ASV that met the filtering criteria and was present in the room was a 100% match to *P. mesoamericanus* with 1070 reads. The other three ASVs matched to *Bos taurus* (Table S1). Only one ASV with a 100% match to *D. rotundus* was present in one control in plate 9 (Table S1). The final plate only yielded sequence data in one control. Only one of the two ASVs present met filtering criteria (100% to *P. mesoamericanus*). The other had a 97% match to *Uroderma convexum* and was removed (Table S1).

Table S1. Breakdown of detections in the negative PCRs for each plate in the 16S region.

| PLATE # | SAMPLING EVENTS | PCR REPLICATE | SPECIES | READ COUNT |
|---|---|---|---|---|
| **1** | A24N & A25D | CLEAN | | |
| **2** | A25N & A26D | 3 | *Artibeus sp.* | 2 |
| **3** | A26N & A27D | 3 | *Desmodus rotundus* | 8 |
| | | 3 | *Alouatta palliata* | 2 |
| **4** | A27N & A28D | CLEAN | | |
| **5** | A28N & A29D | 1 | *Artibeus sp.* | 3 |
| | | | *Desmodus rotundus* | 5 |
| | | | *Pteronotus mesoamericanus* | 3 |
| | | 3 | *Artibeus sp.* | 2 |
| | | | *Cavia porcellus* | 18 |
| | | | *Desmodus rotundus* | 3 |
| **6** | A29N & A30D | CLEAN | | |
| **7** | A30N & M1D | 3 | *Bos taurus* | 5674 |
| | | | *Carollia prespicllillata* | 1430 |
| | | | *Pteronotus mesoamericanus* | 1070 |
| **8** | M1N & M2D | CLEAN | | |
| **9** | M2N & M3D | 2 | *Desmodus rotundus* | 3 |
| **10** | M3N & M4D | 2 | *Pteronotus mesoamericanus* | 10 |
| | | | *Uroderma convexum* | 13 |

**Negative lab controls in COI data**

In the COI region, plate 10 was the only one that yielded no sequence data in the negatives. In plate 1, two of the negatives yielded no sequence data and the sequence data in the third was match to a single ASV with a 99.01% match to *Desmodus rotundus* (Table S2). As it did not meet our filtering criteria, it was removed. The sequence data in the second plate was associated with two ASVs (one in replicate 1 and one in replicate 3). In replicate one, it was a 100% match to *Cavia porcellus* which was used as a positive control, thus this ASV was removed. In replicate 3, it was a 100% match to *D. rotundus* (2 reads) a species that was present in the room and meets the filter requitements. In plate 3, two of the three negatives yielded no sequence data. This data matched to four ASVs all with a 100% match. The species associated with these ASVs are

*Artibeus lituratus*, *D. rotundus*, *Molossus rufus* and *Strunira parvidens.* All these species were present in the room. While plate 4 also had two clean negatives, the third containing matches to seven ASVs making it the most contaminated plate. Two of these ASVs were non-bats (*Phalacrocorax brasilianus*: 13 reads, and *Bos taurus*: 29 reads) and were removed. The other five were bat species. *Dermanura phaeotis* (35 reads), *D. rotundus* (6 reads), *Molossus rufus* (reported as *Molossus sp.,* 244 reads) and *S. parvidens* (17 reads) were present in the room and detected as 100% matches (Table S2). One ASV was a 99.5% match to *Myotis elegans* (11) and was removed as it did not meet filtering criteria. Only one of the negatives did not yield sequence data in the fifth plate. One of the negatives yielded data associated with three ASVs matched to *A. lituratus*, multiple *Molossus sp.* and *Pteronotus fulvus.* As the *Molossus sp.* detections were a 99.5% match to multiple species this ASV was removed. The other contaminated negative had sequence data two ASVs matched to *D. rotundus C. porcellus* (removed as it was the positive control). Plate 6 also only had one negative free from sequence data. The sequence data on the other two were matched to four ASVs. These matched to *D. phaeotis, D. rotundus, Eptesicus furnalis* and *C. porcellus. D. pheaotis* was only a 99.50% match and was removed along with *C. porcellus.* All three negatives yield sequence data in plate 7, matched to only three ASVs. Both *A. lituratus* and *D. rotundus* were 100% matches while the third ASV was a 99.5% match to multiple *Molossus sp.* species (removed). Both plate 8 and 9 only had one contaminated negative. Plate 8 only had one ASV amplified in a single replicate. As it was a 100% match to *Canis sp.* it was removed. In plate 9, there were five ASVs (*D. rotundus*, *Lophostoma brasiliense*, *Molossus sp.* (multiple species), *B. taurus*, *P. brasilianus).* The two non-bat matches (*B. taurus* and *P. brasilianus*) were removed, and the bat detections were associated with 4 or less reads (Table S2).

Table S2. Breakdown of detections in the negative PCRs for each plate in the COI region.

| PLATE # | SAMPLING EVENTS | PCR REPLICATE | SPECIES | READ COUNT |
|---|---|---|---|---|
| **1** | A24N & A25D | 2 | *Desmodus rotundus* | 1 |
| **2** | A25N & A26D | 1 | *Cavia procellus* | 4 |
| | | 3 | *Desmodus rotundus* | 2 |
| **3** | A26N & A27D | 3 | *Artibeus sp.* | 18 |
| | | | *Desmodus rotundus* | 12 |
| | | | *Molossus sp.* | 7 |
| | | | *Sturnira parvidens* | 11 |
| **4** | A27N & A28D | 2 | *Bos taurus* | 29 |
| | | | *Dermanura phaeotis* | 35 |
| | | | *Desmodus rotundus* | 6 |
| | | | *Molossus sp.* | 244 |
| | | | *Myotis elegans* | 11 |
| | | | *Phalacrocorax brasilianus* | 13 |
| | | | *Sturnira parvidens* | 17 |
| **5** | A28N & A29D | 1 | *Artibeus sp.* | 9 |
| | | | *Molossus sp.* | 11 |
| | | | *Pteronotus fulvus* | 2 |
| | | 3 | *Cavia porcellus* | 29 |
| | | | *Desmodus rotundus* | 7 |
| **6** | A29N & A30D | 1 | *Desmodus rotundus* | 5 |
| | | | *Eptesicus furinalis* | 4 |
| | | 3 | *Cavia procellus* | 3 |
| | | | *Dermanura phaeotis* | 3 |
| **7** | A30N & M1D | 1 | *Artibeus sp.* | 3 |
| | | | *Desmodus rotundus* | 5 |
| | | 2 | *Desmodus rotundus* | 5 |
| | | | *Molossus sp.* | 5 |
| | | 3 | *Desmodus rotundus* | 3 |
| | | | *Molossus sp.* | 13 |
| **8** | M1N & M2D | 1 | *Canis sp.* | 2 |
| **9** | M2N & M3D | 1 | *Lophostoma brasiliense* | 2 |
| | | | *Molossus sp.* | 2 |
| | | 2 | *Bos taurus* | 11 |
| | | | *Desmodus rotundus* | 4 |
| | | | *Phalacrocorax brasilianus* | 3 |
| **10** | M3N & M4D | CLEAN | | |

**False positives in 16S data**

There were 13 ASVs that matched to bat species not found or known to be present in the area in the 16S data. Of those 13, five had read counts over 500. The ASV with the highest read count (4192) was a 100% match to the Egyptian fruit bat (*Rousettus aegyptiacus*). This detection is likely a result of secondary transfer on equipment that cannot be deep cleaned (e.g., head lamps, field clothes etc.). Two of the researchers on the research team in Belize work with a captive colony of *R. aegyptiacus*, which is likely why we detected it with such a high read count despite this species only occurring in parts of Africa, the Middle East, Mediterranean and Indian Subcontinent. The second highest read count (1501) was associated with *Lophostoma slivicolum* (100% match). The range of this species only extends to Honduras and thus it is not possible for this detection to be a true positive for the area. However, the next match for this ASV is a 97.78% match to *Lophostoma evotis* which does occur in Belize and was in the field lab during sampling. Thus, it is likely that these two species have similar reference sequences, and this detection is actually associated with *L. evotis*. The next two ASVs were associated *Rhinophylla pumillo* (981 reads, 100% match) and *Nyctinomops macrotis* (925 reads, 100% match). In the case of *R. pumillo*, the genus is only present in south America. There are only two records in the reference database, one of which we produced under an unusual molecular protocol. We exclude this until more is known about variation in the sequenced region for this taxon. While not present in the room during sampling, it is possible that *Nyctinomops macrotis* could be in the area in which case its detection could be a result of drift through the windows and open door. *Nyctinomops macrotis* is also part of the *Molossidae* family which can be difficult to differentiate using DNA and thus this detection could be of one of the *Molossus sp.* present in the room. The final ASV was a 100% match to *Chiroderma trinitatum* (616 reads). While this species is not found in Belize, its sister species *Chiroderma villosum* is present in the Belize and this detection could be a result of drift.

**False positives in COI data**

In the COI data, 12 ASVs matched to bat species not found or known to be in the area. Six of those ASVs were associated with read counts higher than 500. The ASV with the highest read count (10238) was a 100% match to *Anoura geffroyi*. While this species does not occur in Belize, researchers on the trip had been working with these bats prior to coming to Belize and thus their equipment is the likely source of this detection. The ASV matched to *Rousettus aegyptiacus* (599 reads) is also likely to a result of secondary transfer (see false positives in the 16S data). The next highest ASV was a 100% match to *Vampyrodes caraccioli* (2573 reads). While this species does occur in southern Belize it has yet to be confirmed in the study area. As with the 16S data, the detection of *Nyctinomops macrotis* (1458 reads, 100% match) could be a result of drift or a detection of one of the *Molossus sp.* present in the room. It is similar with the ASV matched to *Lophostoma silivcola* (709 reads)*. L. silivola* is not in the area but *L. evotis* which is the next highest match is. Finally, the last ASV is a 100% match to *Diphylla ecaudata* (550 reads). The range of this species does include Belize, and we have previously detected this species in a nearby roost. It is suspected to be present in the study area.

**Appendix 3:** Diversity indices

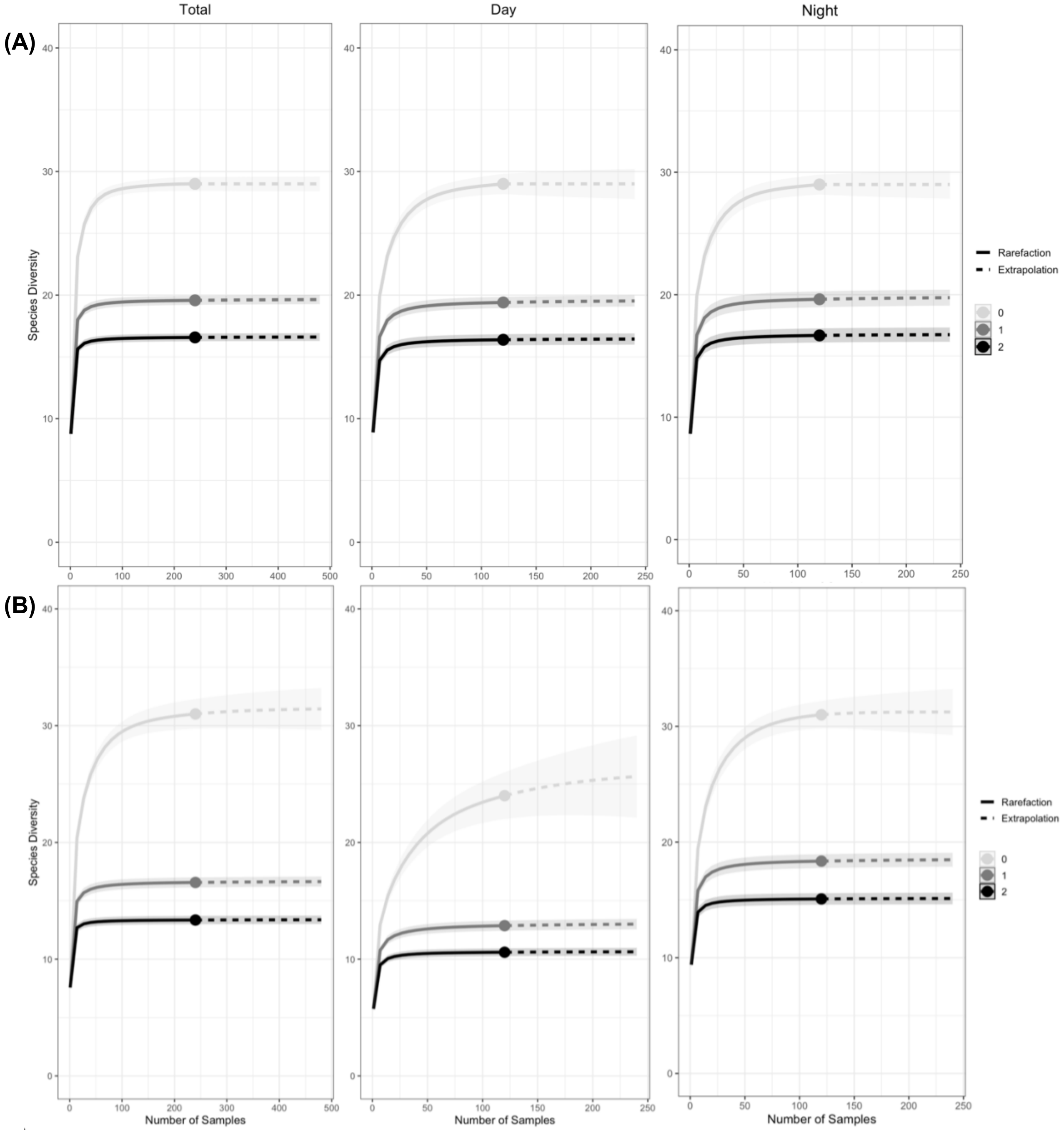


Figure S1. Accumulation curves for bat diversity detected using the 16S (A) and COI (B) markers in the field lab (light grey, q = 0), the Shannon index (dark grey, q = 1) and the Simpson index (black, q = 2). Estimates include 84% confidence intervals and are extrapolated to double the sample value (solid circle). Total coverage (n = 228) represents the complete set of 10 day plus

9-night sampling periods (12 samplers running for 10 days and 9 nights). Note: x-axis is number of samples taken not number of samplers deployed.

**Appendix 4:** Detection Thresholds.

To compare sampling coverage between what we detected and what was confirmed in the room we generated accumulation curves, using the R package iNEXT (Chao et al., 2014; Hsieh et al., 2024), for each night sampled using a detection matrix only of bats that were confirmed to be in the room which in some cases included species that were not detected at all during the sampling period. Then, using the wilcox.test, we ran a series of Wilcoxon tests for each region comparing the sampling threshold of 85%, 95% and total species richness (the total number of species detected).

Using the 16S region, there was a statistically significant difference in species richness ($p = 0.00037$, $W = 0$) and in the 95% threshold ($p = 0.014$, $W = 12.5$; Figure S2) between the dataset encompassing all species detections and the dataset with only the species confirmed to be present that night of sampling, but no difference in the 85% threshold. Using the COI region, there was also statistically significant difference in the species richness ($p = 0.041$, $W = 17$) between the two datasets. However, unlike the 16S data, there was a statistically significant difference for the 85% threshold ($p = 0.0026$, $W = 74$) (Figure S2) but not the 95% threshold.

The statistical difference in species richness is not surprising in the total detections data set as there would still be some detections of species whose DNA was still in the room, but the animal was no longer physically present. The difference in the 16S 95% threshold is also likely due to the differences in species richness between the confirmed presence data and total detections data. In other words, the lower species richness in the confirmed presence data means less samples are needed to detect 95% of the richness. For the COI region, there were some species present that were detectable but were never detected or were not detected on the night they were present (i.e., detected the next day or night). This could explain why the confirmed presence data results in a higher number of samples needed to reach 85% but not 95%. In the confirmed presence data, 85% of the species represents more species than in the total detections data. As more samples are taken, fewer species remain until the only species left to detect in the confirmed presence data are those that were only detected after the animal was no longer physically present. In the total detections data, the 85% threshold is reached much sooner because species can be detected once they have left the room, however the last few rare species are the same in both datasets making the 95% threshold similar regardless of which data set is used.

Table S3. The proportion of species richness detected in 1, 5, 10 and 20 samples for each sampling event and marker.

| | ***16S*** | | | | **COI** | | | |
|---|---|---|---|---|---|---|---|---|
| ***Sampling event*** | ***1*** | ***5*** | ***10*** | ***20*** | ***1*** | ***5*** | ***10*** | ***20*** |
| *A24N* | 0.468 | 0.890 | 0.961 | 0.993 | 0.625 | 0.917 | 0.961 | 0.978 |
| *A25D* | 0.438 | 0.869 | 0.929 | 0.975 | 0.503 | 0.872 | 0.939 | 0.966 |
| *A25N* | 0.520 | 0.925 | 0.978 | 0.991 | 0.805 | 0.952 | 0.974 | 0.992 |
| *A26D* | 0.545 | 0.919 | 0.978 | 0.996 | 0.687 | 0.921 | 0.982 | 0.999 |
| *A26N* | 0.594 | 0.926 | 0.960 | 0.983 | 0.869 | 0.981 | 0.997 | - |
| *A27D* | 0.610 | 0.915 | 0.958 | 0.992 | 0.751 | 0.974 | - | - |
| *A27N* | 0.441 | 0.871 | 0.939 | 0.983 | 0.794 | 0.969 | 0.986 | 0.997 |
| *A28D* | 0.638 | 0.941 | 0.981 | 0.992 | 0.733 | 0.965 | - | - |
| *A28N* | 0.613 | 0.908 | 0.958 | 0.995 | 0.838 | 0.959 | 0.967 | 0.980 |
| *A29D* | 0.586 | 0.906 | 0.952 | 0.976 | 0.629 | 0.948 | - | 0.994 |
| *A29N* | 0.540 | 0.949 | - | 0.996 | 0.699 | 0.922 | 0.950 | 0.972 |
| *A30D* | 0.448 | 0.853 | 0.936 | 0.988 | 0.653 | 0.968 | - | 0.994 |
| *A30N* | 0.540 | 0.919 | 0.958 | 0.982 | 0.466 | 0.911 | 0.965 | 0.985 |
| *M1D* | 0.597 | 0.940 | 0.974 | 0.985 | 0.590 | 0.891 | 0.970 | 0.999 |
| *M1N* | 0.491 | 0.892 | 0.955 | 0.987 | 0.813 | 0.969 | 0.979 | 0.991 |
| *M2D* | 0.569 | 0.924 | 0.977 | 0.998 | 0.420 | 0.813 | 0.901 | 0.945 |
| *M3D* | 0.528 | 0.904 | 0.962 | 0.993 | 0.639 | 0.922 | 0.990 | - |
| *M3N* | 0.511 | 0.926 | - | 0.970 | 0.757 | 0.923 | 0.958 | 0.992 |
| *M4D* | 0.467 | 0.925 | 0.982 | 1.000 | 0.670 | 0.917 | 0.962 | 0.984 |

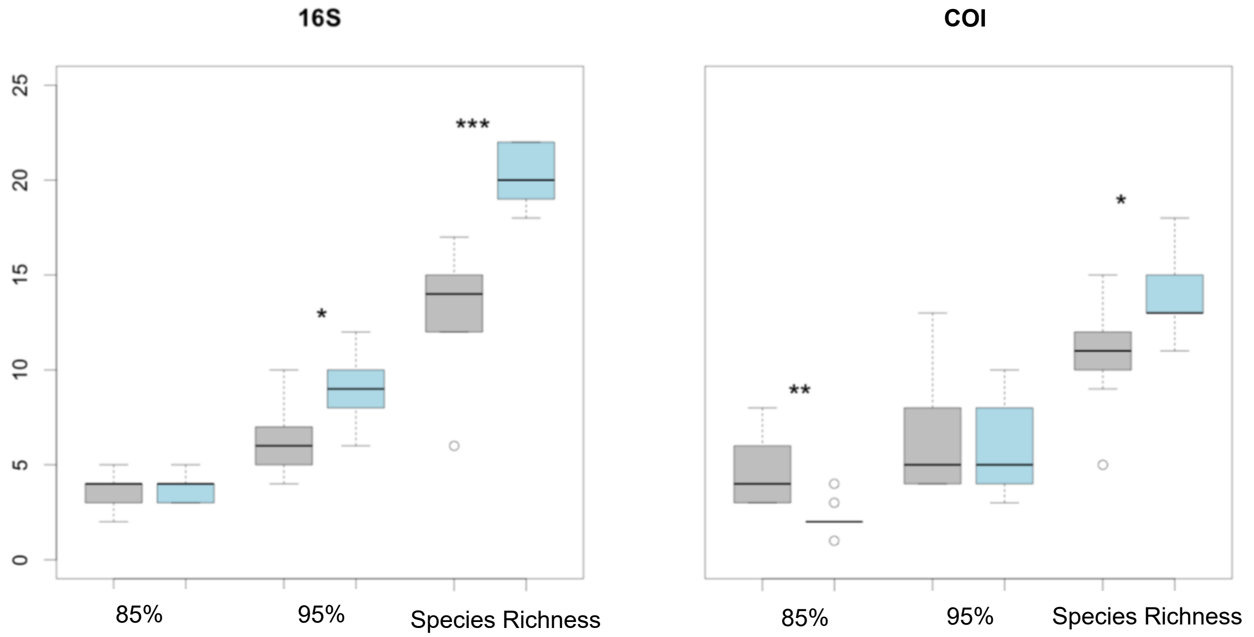


Figure S2. Boxplots comparing species richness, the 85% threshold and the 95% threshold between the confirmed presence data set (grey) and the total detections data set (blue) for each night sampling event. Significance of the Wilcoxon test between the two datasets is as follows: *p<0.05, ** p<0.01, *** p<0.001.
